# Adaptive Cell Size, Merging, Tilting, and Layering in Honeybee Comb Construction

**DOI:** 10.1101/2024.05.29.596484

**Authors:** Golnar Gharooni-Fard, Chethan Kavaraganahalli Prasanna, Orit Peleg, Francisco López Jiménez

## Abstract

Honeybees are renowned for their skills in building intricate and adaptive hives that display notable variation in cell size. However, the extent of their adaptability in constructing honeycombs with varied cell sizes has not been investigated thoroughly. We use 3D-printing and X-ray Microscopy to quantify honeybees’ capacity in adjusting the comb to different initial conditions. Using the average area of natural worker cells as a reference, our findings suggest three distinct construction modes when faced with foundations of varying cell sizes. For smaller cell size, bees occasionally merge cells to compensate for the reduced space. However, for larger cell sizes, the hive uses adaptive strategies like tilting for cells up to twice the reference size, and layering for cells that are three times larger than the reference cell. Our findings shed light on honey-bees’ adaptive comb construction strategies with potential to find applications in additive manufacturing, bio-inspired materials, and entomology.

## Introduction

Honeybee nests are the emergent outcome of the collaborative efforts of thousands of individual bees. These structures play a vital role in the survival and functioning of a bee colony by providing storage space for nectar and pollen, a nursery for brood development, and a structurally stable environment for the various interactions among the nestmates (*1, 2*). The hexagonal structure of the honeycomb, built with a near-optimal minimization of the wax-to-storage space ratio (*3–5*), has fascinated scientists for millennia (*6–8*). This optimization is necessary because wax production incurs a high energy cost (*9*). In particular, to secrete just 1 *lb* (454 *g*) of wax, honeybees need to consume about 8 *lbs* (3.8 *kg*) of honey (*10*).

However, it has been shown that feral bees construct their nests on a range of surfaces such as tree branches, and pre-existing tree cavities, which do not necessarily allow for perfect hexagonal lattices (*11*). Additionally, workers often initiate the process of comb building from multiple locations within the hive simultaneously (*12*). To achieve a coherent structure, the separate pieces of comb need to be merged, and workers must reconcile the differing combination of cell sizes, alignments, etc. at the boundaries, while conforming to the space and resources available (*13*). The resulting structure inevitably contains non-regular hexagons and topological defects with various cell sizes, to adapt to environmental constraints (*14–17*). Adjusting the sizes of hexagons during honeycomb construction may serve additional purposes, often dictated by the colony’s requirements. The need for these adaptations may arise, for instance, when the colony requires space for nurturing workers or drones, or for storing honey, among other reasons (*14*). Despite the frequent occurrence of these size modifications, the degree of adaptability and variability in the structure of the honeycomb in 3D has not been thoroughly investigated.

More than four decades ago, H. R. Hepburn (*10*) studied the comb construction by African Honeybees *Apis melifera adansonii*, by providing them with sheets of beeswax foundations having a range of cell sizes from 170 to 1022 *cells/dm*^2^ (between 1 to 6 times the average size of cells built by bees with no constrains). Using 2D images of the built frames, the study reported various construction solutions and measured working tolerances from the comb built on the foundations with variable cell sizes. Yet, honeycomb represents a three-dimensional structure, and the details and intricacies of it cannot be fully discerned and quantified using two-dimensional images. In this work, we study the adaptability of the comb construction by European honeybees *Apis melifera* L using 3D visualization. We leverage 3D-printing to design repeatable experiments with precisely controlled and carefully quantified cell areas as starter sheets. Our findings corroborate and expand upon prior research by delineating the distinct construction modes employed by bees in response to various initial conditions regarding cell sizes. Furthermore, to highlight the details of each construction mode, we use X-ray microscopy (XRM) to quantify and visualize these intricate structures. Recently, Franklin et al (*18*) have used XRM technology to describe a step-by-step process by which honeycomb is constructed in natural settings which sheds light on the intricate structure of honeycomb cells as they are being built in the absence of a foundation. However, our focus is on identifying various methods of constructing comb pieces, reflecting bees’ ability to adapt to different enforced cell sizes. To that end, our method pairs XRM with 3D-printing, to bring out these various construction modes and their complexity and adaptability to specific initial conditions with variable cell sizes.

To distinguish various foundations, we define the cell size variable (*S*) as a multiple of the inner area of the average honeybee worker cell, which is 12.5 *mm*^2^. This value is derived from the value of the diagonal of a perfect hexagon, which we measured to be 4.4 *mm* on a free comb built with no foundation. Fig. 1A shows a schematic of the shapes of the cells built on the plastic base with *S* = 1, equal to the size of an average worker cell. Fig. 1B shows two snapshots of a 3D-printed frame with *S* = 1 at the start and end of the comb building process. Fig. 1C shows the 3D-visualization of the structure of the comb using X-ray data from a section of the frame shown in panel B. The details of how we process the raw X-ray data to generate 3D-visualizations of the comb structure are described in the Methods Section . Using this size as our baseline, we supply the hives with custom 3D-printed frames with regular hexagonal patterns of various cell sizes, ranging from half to four times the size of a reference cell (*i.e.*, 0.5 *≤ S ≤* 4). Our goal with changing the foundation’s cell sizes is to examine the bees’ capacity to adjust the comb structure to consistently smaller or larger cells. Fig. 1D provides an overview of our results highlighting different construction modes observed on our 3D-printed frames. It includes photographs of these building modes, followed by schematics for adaptive strategies used for honeycomb construction with varying cell sizes. The details of the structure of the honeycomb in each mode is discussed in the following section.

## Results

We begin this section by discussing the different construction modes identified in our experimental data, followed by a more detailed description and visualization of the overall structure of the comb in each category. We specifically identify three different building modes—merging, tilting, and layering—relative to the given foundation sizes. The intricate properties of each construction category, along with 3D-visualization using XRM, is presented in the following sections.

### Merging

To examine how bees adapt their comb structure when faced with smaller cell sizes, we manipulate the cell areas of the printed patterns, reducing the area of the 3D-printed cells to 75% the average worker cell (*i.e. S* = 0.75). Fig. 2 A-D illustrates the evolution of the comb construction on one of these experimental frames over a 20-day period. There is an apparent pattern at work from the early stages of comb construction, indicating the swarms’ consistent recognition of and adaptation to this new cell size pattern. As shown in the insets of Fig. 2A-B, even though most of the initially drawn hexagons start from the plastic edges on the printed foundation, some cell walls are built with an angle that will eventually merge with the walls from the adjacent cells built with a reverse tilt. This pattern repeats occasionally until the whole frame is covered with hexagonal cells. The term “merging” describes the occasional combination of several small cells to create the necessary space to construct natural worker-size hexagons.

**Figure 1:**
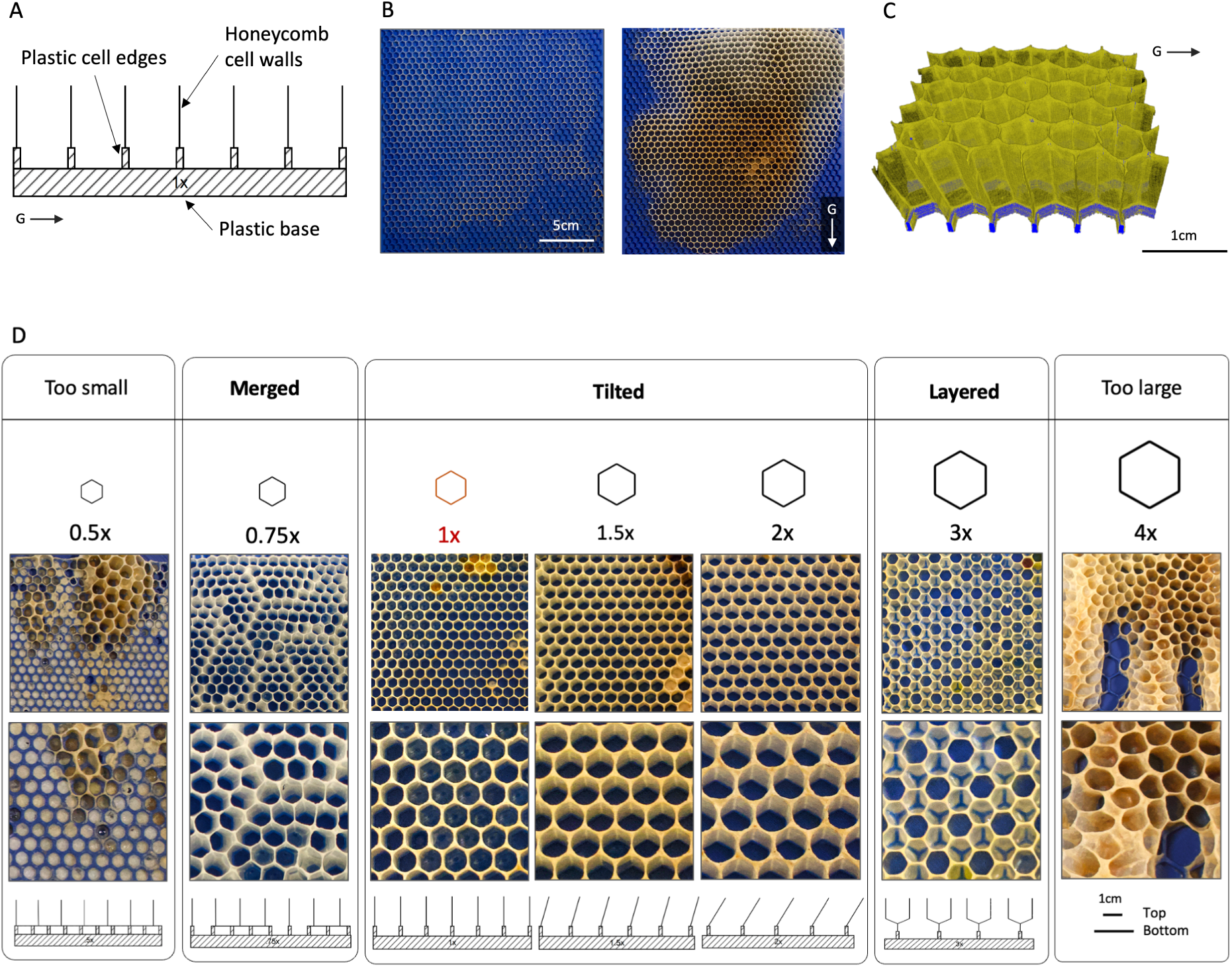
A) Schematic of the honeycomb cells built on a control frame with *S* = 1. B) Two snapshots of the initial frame with the plastic, 3D-printed foundation along with the same frame after 20 days. The printed cell size on this frame is *S* = 1. C) XRM-based 3D-visualization of a section of the frame with *S* = 1, segmented to highlight the comb in dark yellow and the plastic in blue. The direction of gravity is indicated by the arrow labelled *G*. D) Overview of the different modes of honeycomb construction on 3D-printed frames with varying cell sizes. The top row shows the building mode in relation to the cell sizes on each foundation, displayed beneath them. The subsequent row presents images of the constructed honeycomb on the foundations, accompanied by a zoomed-in view beneath. The bottom row illustrates schematics depicting adaptive strategies for honeycomb construction with differing cell sizes.

Fig. 2D shows the frame after the comb construction is complete, but prior to the filling of cells with honey or brood, at which point X-ray imaging is conducted. The series of images in Fig. 2E-H demonstrate the X-ray results. In particular, Fig. 2E and Supplementary Movie S1 illustrates a 3D-reconstruction of a 5 *cm ×* 5 *cm* section of the frame in Fig. 2D, highlighting plastic in blue and comb in dark yellow. To better visualize the merging process, the cross-sections of the X-ray are presented in Fig. 2F-H, using planes containing two of the Cartesian axis, shown in green (XZ), blue (YZ), and red (XY) in Fig. 2E. For instance, the positioning of the cell walls on the XZ slice shown in Fig. 2F highlights how comb material is used to combine adjacent plastic edges beneath the cells constructed on top of them.

Additionally, in the bottom view of the sample in Fig. 2I, it can be observed that some of the printed cells are entirely covered with comb. This coverage aims to provide additional space for the initiation of a normally-sized hexagonal lattice. The distribution of covered cells on a larger section of the frame can be viewed in Fig. 2J, with covered cells shown in gray. Interestingly, we find that on average, one out of every four cells built on the frames with *S* = 0.75 foundation is covered. This ratio suggests that bees employ the “merging” strategy to effectively transform a slightly small hexagonal grid into their commonly-used size. In other words, they combine one out of four cells to compensate for the 25% less space imposed by our 3D-printed foundations with *S* = 0.75.

**Figure 2:**
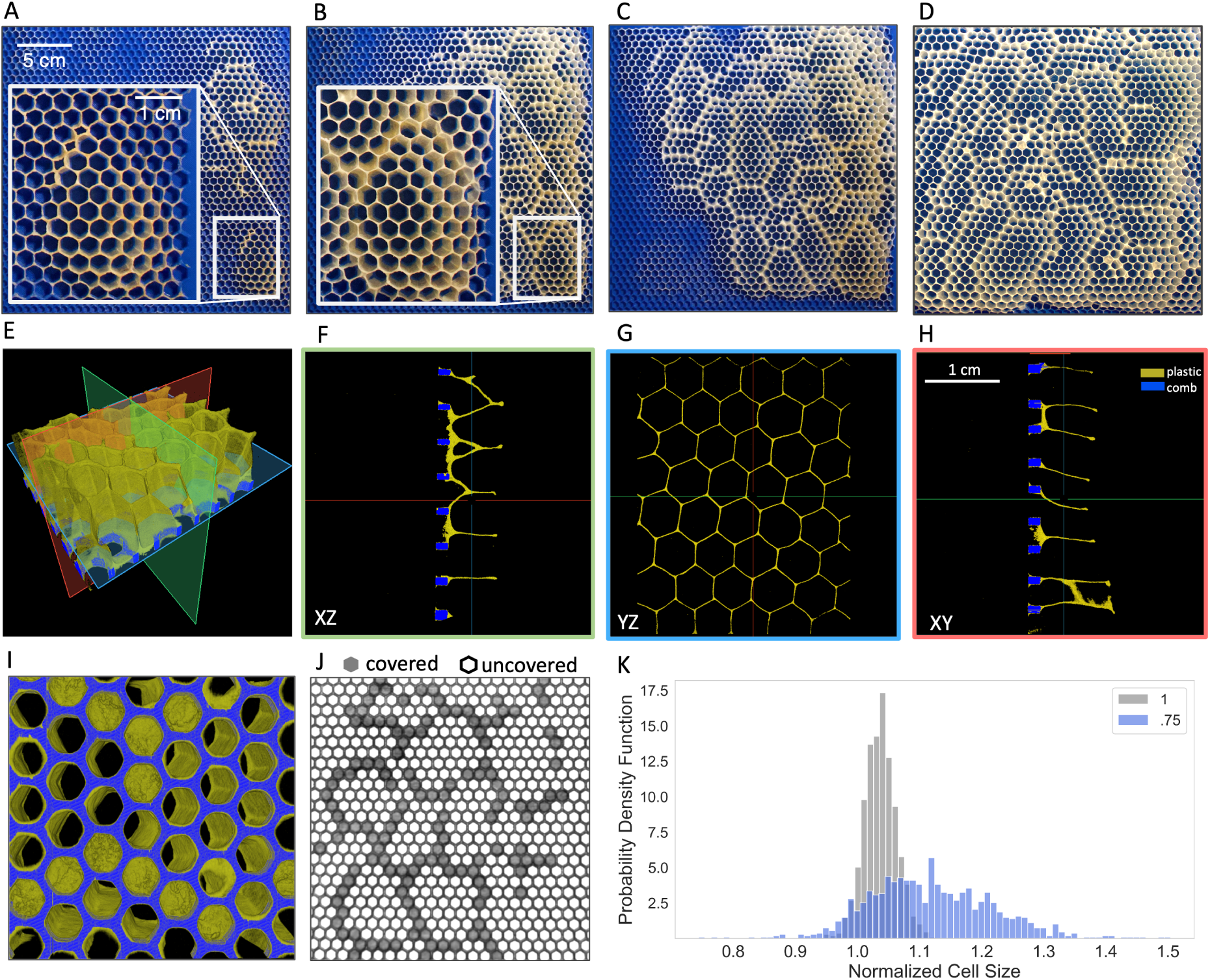
A) The 3D-printed frame of cells with *S* = 0.75 thinly coated with beeswax after 3 days of being in the hive. The inset shows the zoomed in section of the constructed comb highlighting cells with tilted walls. B) The same frame after a week. The inset shows the same zoomed in section as in panel A, demonstrating how the adjacent cells with tilted walls will be merged once they are fully constructed, effectively concealing the smaller cell below on the foundation. C) The same frame after 10 days, and D) after 20 days. E) 3D visualization of a 5 *cm ×* 5 *cm* section of the comb. Three planes shown in green, blue and red are used to create cross-sections of the comb structure, shown in F-H). In all of these slices the comb is shown in dark yellow while the plastic is shown in blue. I) The bottom view of the 3D reconstructed volume showing the cells that are covered with comb and the uncovered cells. J) 2D visualization of a larger section of the same sample with uncovered cells in white and covered cells in gray. K) Comparison of the probability density function of cell sizes built with *S* = 0.75, shown in blue, with the reference cells, shown in gray.

Remarkably, while some of the printed cells are filled with wax, the top view of the comb structure, shown in Fig. 2G, still features a regular hexagonal grid with cell sizes that are comparable to the average worker cell size. For comparison, Fig. 2K depicts the distribution of cell sizes for cells built on the frames with *S* = 0.75 in blue, along with the distribution of the reference cell sizes shown in gray. The blue distribution covers a similar region but is considerably wider (*S* = 1.04 *±* 0.02 versus *S* = 1.12 *±* 0.11), suggesting less consistency in the cell sizes of the comb built on the foundation with *S* = 0.75. The higher standard deviation could be attributed to the discrepancy between the given and desired cell sizes and the inherent randomness of the merging process. It is worth noting that the merging behavior during honeycomb construction is not specific to these experimental frames with smaller given foundations. Similar pattern can sometimes be observed on the comb built on some of the commercially available frames with worker-size foundations. See Fig. S3 in the Supplementary Materials for an image of the comb built on one of the commercial frames inside our hives using the merging strategy. This adjustment may be made to meet the colony’s specific needs for constructing larger cells, such as those used for raising drones.

### Tilting

The next mode of construction corresponds to the case with the given foundation of cells with 1 *≤ S ≤* 2, where bees use the provided patterns but increase the tilt angle of the cells as a function of the cell size. Fig. 3A presents a series of 2D-images of the comb built on frames with *S* = 1, 1.5, and 2. The corresponding 3D-reconstructions of the comb built on these frames are presented in Fig. 3B, which are the result of scanning sections of the combs highlighted with red boxes in Fig. 3A. Tracing the plastic edges shown in blue and the comb in dark yellow across the three samples displayed in Fig. 3B reveals a consistent behavior: bees construct honeycomb cells along the edges of plastic foundations, regardless of the increasing cell sizes observed in the samples. The distribution of the built cell sizes on frames with foundations ranging from 1 to 2, as depicted in Fig. 3C, demonstrates a monotonic increasing trend indicating a clear positive correlation between the foundation and the final cell size. This confirms the bees’ ability to effectively utilize the provided patterns by adjusting honeycomb cell sizes to match the given foundations. According to our observations, these larger cells that are too large for raising workers, are either used for raising drones or storing honey (see Fig. S4 for some examples). For context, the dashed line in Fig.3C marks the average cell size of natural drone brood cells. While the dashed line aligns well with the *S* = 1.75 and closely follows the *S* = 1.5 distributions, it is quite far from *S* = 2. Nevertheless, it is interesting to observe that in our experiments all of the larger cell types are utilized for drone rearing when required.

Additionally, we notice that the bees notably increase the tilt as the foundation’s cell sizes increase. Previous studies exploring the natural tilt of the honeycomb cells have indicated that a standard worker cell tilts up to 13*^◦^* (*15, 19, 20*). It has also been established that the direction of this tilt is influenced by gravity, meaning that bees consistently tilt the cells upward (*21–23*), against the direction of gravity. In our experiments, we note that cells on the control frames with *S* = 1 are tilted, on average, by 8.75*^◦^* against the direction of gravity. Table 1 shows the average tilt of the comb cells built on various foundation sizes. It is evident that there is a gradual rise in cell tilts with increasing values of *S*. However, this relationship is not strictly linear. Notably, cells built on foundations with *S* = 1.75 are, on average, smaller than those with *S* = 1.5 and they also exhibit a wider range of tilts (with a standard deviation of 2.73*^◦^*). This trend suggests a non-linear relationship between cell size and tilt, indicating that tilt may be influenced by various hive structural factors, such as cell position, usage, content, etc. For comparison, the last row in Table 1 shows the tilt angle distribution for natural drone cells.

Prior studies have suggested various hypotheses to explain the natural tilt of the honeycomb cells. Mullenhoff (*24*) posited that this tilt prevents honey outflow. However, a recent study by Oeder et al. (*20*) challenges this idea, proposing instead that the tilt redistributes weight onto the midwall, potentially increasing the comb’s carrying capacity. In our experiments, we find that the amount of tilt effectively reduces the cross-sectional area of the cell, when considered perpendicular to the walls. For example, when considering the case of *S* = 2 and cutting the sample with a plane perpendicular to the walls at the top of the sample, the cross-sectional area of the hexagons at that layer is, on average, 0.77 times the area of the printed hexagons. This means that the cross-sectional area is equivalent to *S* = 1.54, a significant reduction with respect to the provided lattice *S* = 2. See Supplementary Movies S2-6 for the 3D reconstruction of all the samples built with tilting cells.

**Figure 3:**
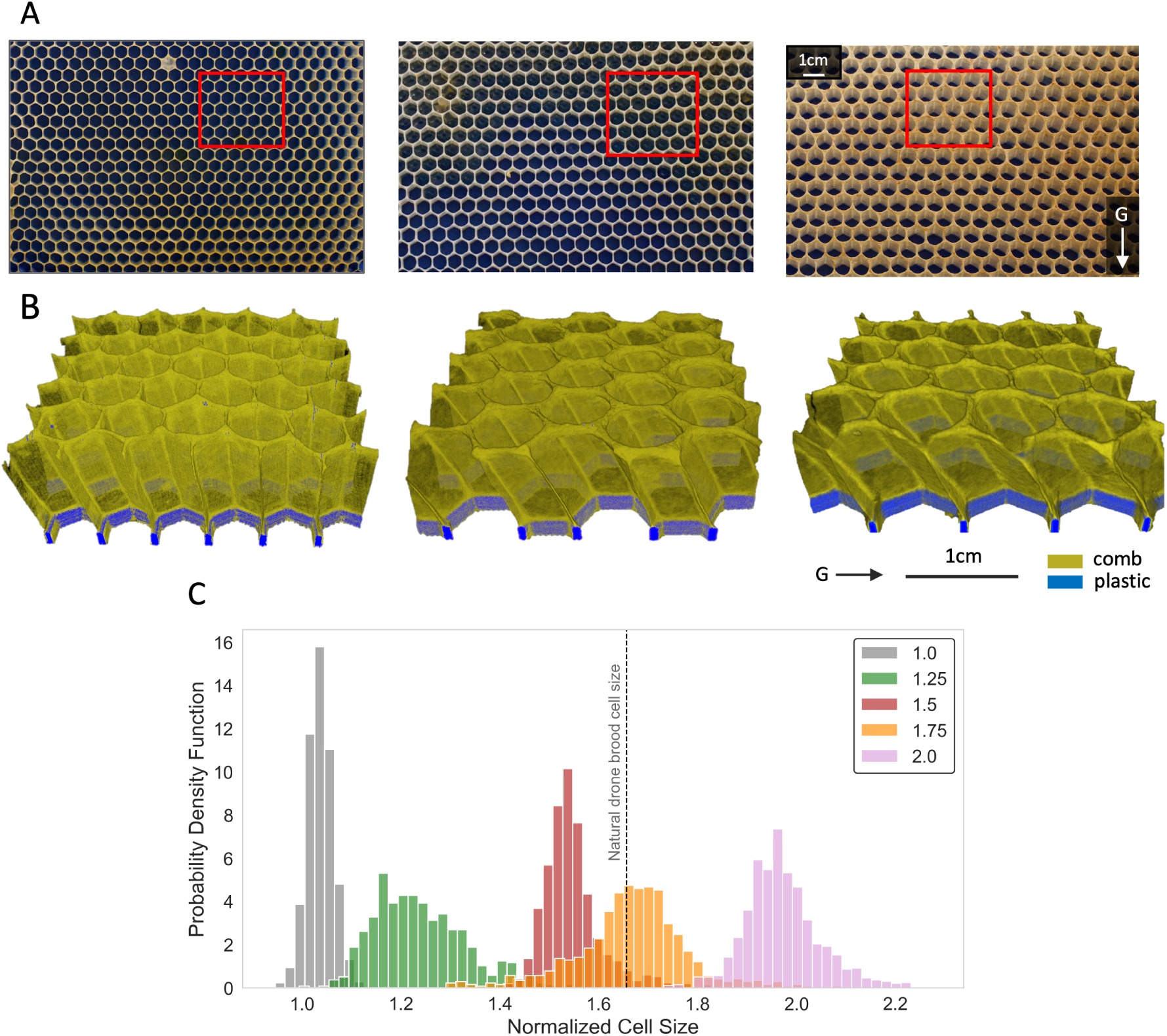
Results of the comb built on samples with 1 *≤ S ≤* 2 foundation. A) The 2D images of the frame with *S* = 1, 1.5, and 2. B) The 3D reconstruction of the sections of the comb, highlighted with a red box in panel A, using X-ray data, segmented to show comb in dark yellow and the plastic in blue. C) Distribution of the final cell sizes for the comb built on the frames with larger foundations. The color shows the value of *S*, and the vertical dashed line marks the average cell size of natural drone brood cells.

**Table 1:** Mean and standard deviation of cell tilts measured across various given cell sizes (S). The tilt for the drone comb is measured manually using images of natural drone comb built on a flat surface, such as Fig. S5.

| Foundation cell size ($S$) | Average normalized cell size | Average tilt |
| --- | --- | --- |
| 1 | 1.04 +/- 0.03 | 8.74 +/- 2.29 |
| 1.25 | 1.24 +/- 0.09 | 9.44 +/- 1.97 |
| 1.5 | 1.55 +/- 0.06 | 22.64 +/- 0.85 |
| 1.75 | 1.67 +/- 0.11 | 16.7 +/- 2.73 |
| 2 | 1.97 +/- 0.08 | 27.1 +/- 1.65 |
| Natural drone comb | 1.61 +/- 0.1 | 20.02 +/- 2.42 |

### Layering

In this section we explore the cells built on frames with areas three times larger than those of worker cells (*i.e. S* = 3), which result in a new mode of construction, see Fig. 4. Contrary to the previous cases, the bees did not adapt the provided pattern by simply constructing larger or more tilted cells. Instead, they use the corners of the 3D-printed cells as the foundation for building a new layer of cells, forming a two-layer structure; see Fig. 4A. This 2D view shows that the cells on the upper layer are centered at the corners and supported by the edges of the 3D-printed hexagons. This arrangement results in an additional cell, located at the center of the provided hexagons, though smaller in size. This cell extends until the base of the 3D-printed frame, merging with the provided hexagon. This structure becomes more clear in the 3D view of the comb, depicted in Fig. 4C and Supplementary Movie S7 that shows how the honeycomb cells (shown in dark yellow) are positioned on top of the plastic edges (shown in blue).

As shown in Fig. 4B, the distribution of final cell sizes is wider (the standard deviation is 0.02 for *S* = 1 vs. 0.07 for *S* = 3), but quite similar to the cell sizes built on the control frames. This suggests that the two-layer strategy employed by bees effectively transforms the given foundation of *S* = 3, which is too large to serve any purpose in the hive, to *S* = 1, which is the most commonly used. Overall, this symmetrical structure is comprised of six new normal-size hexagons superimposed on the three-times larger given hexagon, resulting in the formation of another hexagon at the center. Consequently, as shown in Fig. 4A, each gray hexagon on the top layer derives about 1/3 of its area from the 3D-printed (blue) cell and about 2/3 from the neighboring cells upon which it is constructed. It is worth noting that, while we printed the three-times-larger hexagon with a vertical orientation, with its side walls perpendicular to the top bar, the resulting honeycomb lattice built by bees features a horizontal orientation, with side walls parallel to the top bar. While the former orientation is commonly used in commercially-available foundations and it is the orientation most commonly observed in naturally built comb, the latter has been found to be approximately 30% stronger structurally (*10*).

**Figure 4:**
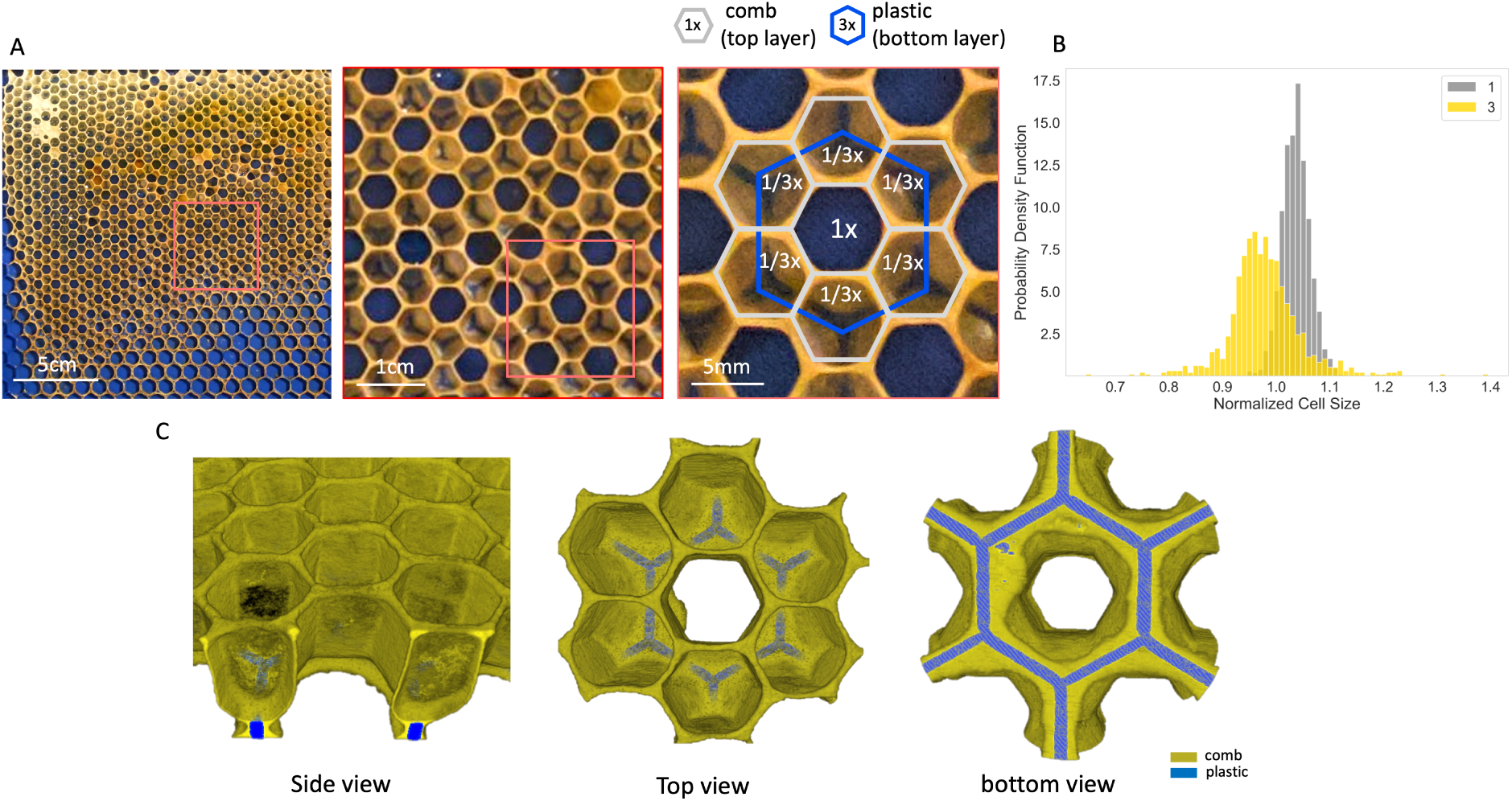
Results of the comb built on a sample with *S* = 3 foundation. A) Image of the comb built on the frame with *S* = 3, followed by the zoomed in view of the area highlighted with a red square. B) Comparison of the distribution of the final cell sizes built on foundations with *S* = 3 in yellow, with the ones built on the reference frame with *S* = 1 in gray. C) Three views of the 3D reconstruction focusing on one given hexagon and the surrounding cells using X-ray data, showing the comb in dark yellow and the plastic in blue.

## Discussion

Within the range of cell sizes we examined, which is 0.5 *≤ S ≤* 4, it became apparent that bees do not effectively utilize the patterns on the provided foundation when *S* = 0.5, or *S* = 4. For *S* = 0.5, the bees either disregard the provided pattern entirely or fill the small cells with wax (see Fig. 1D). This allows them to initiate the comb construction on a flat foundation. This behavior resembles the nesting practices of wild honeybees found in tree cavities with numerous small cracks and crevices (*25*). In such natural settings, bees tend to create a flat and waterproof surface by using tree resins (propolis) to seal and coat the tiny gaps and cracks, effectively preventing the growth of mold and bacteria (*11*). In the case of *S* = 4, the bees are unable to use the given foundations to build a uniform structure. Most of the comb built on this size foundation seem to be randomly constructed in all directions, sometimes growing outwards perpendicular to the frame (see Fig. 1D). We have also collected data using foundations with other sizes such as *S* = 2.5 and *S* = 3.5. On these frames we observed an inconsistent mix of tilted and layered hexagonal cells, please see Fig. S6 on the Supplementary Materials section for examples.

For the remaining foundations tested (0.75 *≤ S ≤* 3) our results show different mechanisms by which honeybees adapt regular hexagonal foundations of different sizes to satisfy their needs. In the cases with *S* = 0.75 and *S* = 3, the provided cell sizes are too different from those naturally used by the bees, so they need to adapt them to build a new lattice. In the case of smaller cells, the size mismatch between both layers is resolved by effectively blocking some of the given cells, so that their space is distributed among their neighbor cells. This can be understood as the discrete version of systems in which geometric mismatch between two layers is resolved through localization of deformation, from cracks in drying films or mud (*26, 27*) to ridges in pre-stretched elastic bilayers (*28, 29*), which result in similar patterns to those created by the blocked hexagonal cells. In the case of *S* = 3, the bees use the plastic edges of the provided frame as a foundation to build a new regular hexagonal lattice. The new hexagons are centered on the edges and vertices of the provided lattice, constructing a new layer of hexagonal lattice with cell size close to *S* = 1. This structure closely resembles the comb construction seen in African honeybees on a foundation documented in (*10*). While Hepburn described this middle cell as a “false” cell that is not easily noticed in the completed comb, our experiments reveal that the broad base of this central cell can serve as an additional storage space for honey and pollen. In contrast, in the range 1 *≤ S ≤* 2, the bees extend the provided lattice, constructing larger cells while increasing the tilt of the cells as their size increases. For a comprehensive comparison of the cell sizes constructed on all provided foundations, please refer to Fig. S7 in the supplementary materials section.

In our previous work (*17*), we introduced a 2D model capable of replicating the patterns and defects on the comb structure arising from various geometric frustrations. With the identification of novel building modes, particularly the two-layered pattern in our current study, now we are motivated to extend our 2D model into a 3D framework. This extension will enable us to delve deeper into understanding the costs and benefits associated with such structural adaptations, which could be leveraged in the design of novel lightweight structures conforming to complex irregular geometries. Furthermore, given the bees’ demonstrated capacity to construct and utilize large tilted cells, our future work involves rationalizing the relationship between cell size and angle, starting with the two reasons already proposed in literature. With that objective in mind, we will examine the mechanical stability of the comb under larger tilt values, as well as the fluid mechanics that govern honey outflow within these structures. These future directions aim to enhance our comprehension of the intricate dynamics underlying comb construction and functionality of these adaptive structures.

## Methods

To collect data about the structure of the comb, we establish a set of behavioral assays combined with X-ray imaging. The focus of this study is on the cell size (0.5 *≤ S ≤* 4) which is varied systematically using 3D-printed panels so that its effect can be isolated and quantified. The design work for preparing the 3D-printed frames is conducted using the SolidWorks 2019 CAD design software. All of our experiments are performed using colonies of European honeybees *Apis melifera* L. at the Peleg lab apiary in Boulder, Colorado. We use langstroth hives with 19 *in* (480 *mm*) frames, with 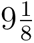 *in* (230 *mm*) depth, and 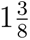 *in* (35 *mm*) width. Each frame consists of a pair of 3D-printed plates of the size 212 *×* 212 *mm* that are placed opposite to each other. For our control set with *S* = 1, each hexagonal element on the 3D-printed plates has a side-length of 2.7 *mm* and outer area of 18.94 *mm*^2^ (including the cell walls). All the other configurations are designed relative to the area of our control set. To confirm the observed modes of construction, we collected at least three repetitions of samples for each cell size, *S*. See the Supplementary Data section of the Supplementary Materials for pictures of these samples. To account for at least three repetitions of each of the configurations under study, we scatter a total of 50 3D-printed experimental frames into ten honeybee colonies and fill the rest of the space in the hives with normal frames with full plastic foundations.

To stimulate comb building, we apply a fine layer of natural beeswax to the printed foundation. The comb construction progress is monitored through regular photography sessions, capturing images of both sides of the experimental frames. To ensure minimal disruption during photography, we set up our photography station within the apiary, allowing us to quickly return the frames to their original locations inside the hives. Once the experimental frames are fully built, they are carefully extracted from the hives, and kept in the lab for further analysis. We obtain high-quality images of the comb using a Nikon DSLR camera and controlled lighting, and a black background. The camera is mounted on a fixed metal structure 2 *m* from the comb to avoid distortion at the edges of the image and to keep a constant pixel-to-millimeter relationship. At this distance and camera resolution, 1 *mm* = 15 pixels.

To further analyze the honeycomb structures obtained after the completion of the experiments, we used 3D X-ray microscopy (XRM) to scan 5 *cm ×* 5 *cm* sections of the comb constructed on our experimental frames. This provides a series of high resolution cross-sections of pieces of the comb that can be combined to generate a 3D virtual model. All tomography scans are performed using *µ − CT* ZEISS Xradia 520 Versa (Carl Zeiss X-ray Microscopy Inc, Pleasanton, CA, USA). The X-ray source is set to voltages in range 80-90 *kV* and power 7 *W*, while the detector camera is set with an exposure time of 2 seconds for each of the 2034 images captured by optical magnification 0.4. Image processing and segmentation on the X-ray data are conducted using Dragonfly (Object Research Systems, Montreal, QC, Canada). See the “X-ray data analysis” section in the Supplementary Materials for a detailed account of our processing pipeline.

## Data Availability

The data used in this study can be found at, https://datadryad.org/stash/dataset/ doi:10.5061/dryad.z8w9ghxmw.

## Code Availability

The code used to analyze the X-ray data in this study can be found at https://github.com/peleg-lab/honeycomb-adaptive-cell-size.

## Acknowledgments

O.P. and F.L.J. acknowledge funding from the University of Colorado Boulder RIO Seed Grant Program, and NSF grant 2210628. We also acknowledge funding from the University of Colorado Boulder, BioFrontiers Institute (internal funds). Any opinion, findings and conclusions or recommendations expressed in this material are those of the authors(s) and do not reflect the views of the NSF. XRM data collection and parts of the analyses were performed at MIMIC facility, at CU Boulder (RRID:SCR 019307). We thank Seneca Kristjonsdottir and Christopher Borke for bee management, Adrian Gestos for XRM training and trouble shooting, Paul Bontempo, Richard Terille, Anna Rahn, and Anna Simone for their assistance with data collection and organization.

## Author Contributions

G.G., F.L.J. and O.P. designed research; G.G. performed research; G.G., C.K.P., F.L.J. and O.P. contributed new reagents/analytic tools; G.G. and C.K.P. analyzed data; G.G., C.K.P., F.L.J. and O.P. wrote the paper.

## References

1. Thomas D Seeley. The wisdom of the hive: the social physiology of honey bee colonies. Harvard University Press, 2009.

2. Paul Siefert, Nastasya Buling, and Bernd Grünewald. Honey bee behaviours within the hive: Insights from long-term video analysis. Plos one, 16(3):1–14, 2021.

3. Thomas C Hales. The honeycomb conjecture. Discrete & Computational Geometry, 25(1):1–22, 2001.

4. Forrest H Kaatz, Adhemar Bultheel, and Takeshi Egami. Order parameters from image analysis: a honeycomb example. Naturwissenschaften, 95(11):1033, 2008.

5. Tim Räz. On the application of the honeycomb conjecture to the bee’s honeycomb. Philosophia Mathematica, 21(3):351–360, 2013.

6. John Hunter. Viii. observations on bees. Philosophical Transactions of the Royal Society of London, (82):128–195, 1792.

7. Charles Darwin. On the origin of species, 1859. London: John Murray, 1859.

8. 8. DA Thompson. On growth and form. Cambridge : University Press ; New York, 1945.

9. Darcy Wentworth Thompson and D’Arcy W Thompson. On growth and form, volume 2. Cambridge university press Cambridge, 1942.

10. HR Hepburn. Comb construction by the african honeybee, apis mellifera adansonii. Journal of the Entomological Society of Southern Africa, 46(1):87–101, 1983.

11. 11. Thomas D Seeley. The lives of bees: the untold story of the honey bee in the wild. Princeton University Press, 2019.

12. Peter R Marting, Benjamin Koger, and Michael L Smith. Manipulating nest architecture reveals three-dimensional building strategies and colony resilience in honeybees. Proceedings of the Royal Society B, 290(1998):20222565, 2023.

13. TD Seeley and RA Morse. The nest of the honey bee (apis mellifera l.). Insectes Sociaux, 23(4):495–512, 1976.

14. HR Hepburn and LA Whiffler. Construction defects define pattern and method in comb building by honeybees. Apidologie, 22(4):381–388, 1991.

15. Ming-Xian Yang, Ken Tan, Sarah E Radloff, Mananya Phiancharoen, and H Randall Hepburn. Comb construction in mixed-species colonies of honeybees, apis cerana and apis mellifera. Journal of Experimental Biology, 213(10):1659–1664, 2010.

16. Michael L Smith, Nils Napp, and Kirstin H Petersen. Imperfect comb construction reveals the architectural abilities of honeybees. Proceedings of the National Academy of Sciences, 118(31), 2021.

17. Golnar Gharooni Fard, Daisy Zhang, Francisco López Jiménez, and Orit Peleg. Crystallography of honeycomb formation under geometric frustration. Proceedings of the National Academy of Sciences, 119(48):e2205043119, 2022.

18. Rahul Franklin, Sridhar Niverty, Brock A Harpur, and Nikhilesh Chawla. Unraveling the mechanisms of the apis mellifera honeycomb construction by 4d x-ray microscopy. Advanced Materials, 34(42):2202361, 2022.

19. H Randall Hepburn. Honeybees and wax: an experimental natural history. Springer Science & Business Media, 2012.

20. Robert Oeder and Dietrich Schwabe. The upward tilt of honeycomb cells increases the carrying capacity of the comb and is not to prevent the outflow of honey. Apidologie, 52:174–185, 2021.

21. Gabriele Von Oelsen and Eva Rademacher. Untersuchungen zum bauverhalten der honigbiene (apis mellifica). Apidologie, 10(2):175–209, 1979.

22. H Martin and M Lindauer. Sinnesphysiologische leistungen beim wabenbau der honigbiene. Zeitschrift für vergleichende Physiologie, 53:372–404, 1966.

23. Shunhua Yang, Shangkao Deng, Haiou Kuang, Danyin Zhou, Xueyang Gong, and Kun Dong. Evaluating and comparing the natural cell structure and dimensions of honey bee comb cells of chinese bee, apis cerana cerana (hymenoptera: Apidae) and italian bee, apis mellifera ligustica (hymenoptera: Apidae). Journal of Insect Science, 21(4):1, 2021.

24. 24. Karl Müllenhoff. Ueber die entstehung der bienenzellen. Emil Strauss, 1883.

25. Thomas D Seeley. Adaptive significance of the age polyethism schedule in honeybee colonies. Behavioral ecology and sociobiology, 11:287–293, 1982.

26. EM Kindle. Some factors affecting the development of mud-cracks. The journal of Geology, 25(2):135–144, 1917.

27. Wai Peng Lee and Alexander F Routh. Why do drying films crack? Langmuir, 20(23):9885–9888, 2004.

28. Jianfeng Zang, Xuanhe Zhao, Yanping Cao, and John W Hutchinson. Localized ridge wrinkling of stiff films on compliant substrates. Journal of the Mechanics and Physics of Solids, 60(7):1265–1279, 2012.

29. Rashed Al-Rashed, Francisco López Jiménez, Joel Marthelot, and Pedro M Reis. Buckling patterns in biaxially pre-stretched bilayer shells: Wrinkles, creases, folds and fracture-like ridges. Soft Matter, 13(43):7969–7978, 2017.

