## Supplementary material for "Adaptive Cell Size, Merging, Tilting, and Layering in Honeybee Comb Construction": SI_Honeycomb_size_paper.pdf

### Supplementary Materials for
### Adaptive Cell Size, Merging, Tilting, and Layering in Honeybee Comb Construction

Golnar Gharooni Fard et al.

This PDF file includes:

- Supplementary Text
- Supplementary Table S1
- Supplementary Figures S1 to S7
- Supplementary Movies S1 to S7
- Supplementary Data

### Supplementary Text

#### X-ray data analysis

X-ray tomography scanning of our experimental sample generates a stack of cross-sectional tomography images which forms a 3D model of the sample when digitally stitched together. Each tomography image comprises of the following elements– air, honeycomb, and the plastic base, each captured with a different pixel intensity depending on the radiodensity of the element. The preprocessed X-ray data for a sample with $S = 1$ is shown in Fig. S1A, with the areas of interest labelled as honeycomb, plastic cell edges and plastic base. We develop an image processing pipeline that converts the stack of tomography images to a form that is more amenable to visualization and analysis. We mainly use computer vision techniques, with the OpenCV [1] library in Python, to conduct all our image analyses. The main objective of our image processing pipeline is to filter out unwanted parts of the raw X-ray data and segment particular regions of interest. This is done in three steps. First, we employ image thresholding to filter out portions of the image containing air, using an in-house heuristic to determine the optimal threshold value for this purpose. Next, we filter out the plastic foundation while maintaining the plastic cell edges, for creating clear 3D visualizations. Here we use a combination of convolution with a specially-designed kernel and morphological operations. The final step is to identify and segment the plastic cell edges. We achieve this by using heuristic-based thresholding, similar to the first step. In addition, we systematically analyze the angle of tilt in the cells using the scientific image analysis software Dragonfly [2]. The following sections describe details of all the steps in our image processing and analysis pipeline.

##### 1. Histogram-based background filtering

To filter out regions containing air in the unprocessed X-ray tomography images, shown in Fig. S1A, we use heuristic-based thresholding. The histogram of pixel intensities for the XZ slice corresponding to the sample in panel A is depicted in Fig. S1B. The prominent gray peak in this histogram highlights the air pixels that constitute approximately 88.9% of the unprocessed tomography images. Our method uses the second derivative of the image histogram to find the optimum threshold value for the effective segmentation of air from data (*i.e.* honeycomb and plastic). The resulting threshold value is highlighted with a red-dashed vertical line in Fig. S1B. It is worth noting that, other methods of background filtering, such as absolute and adaptive thresholding [3, 1] offered sub-optimal results for our objectives. In particular, absolute thresholding requires manual determination of the optimum threshold value for each image in the dataset. Owing to the large number of images in each of our datasets, this is not feasible to do efficiently and is precisely the process that we have automated in our method. In addition, the quality of results produced by adaptive thresholding were inconsistent across images within a given dataset and the amount of noise in these results exceeded our tolerance for error.

#### 2. Segmentation of plastic frame

To segment the plastic from honeycomb, we first identify and mask out the plastic base using kernel convolution and morphological operations [1]. Next, we segment the plastic cell edges imprinted on the plastic base which act as seeds for honeycomb construction. These regions are labelled in Fig. S1A. Masking out the plastic base is done in two stages. Initially, we identify pixels delineating the boundary between the plastic base and air. Upon applying histogram thresholding to the image, we observe that one side of the plastic base boundary consists solely of black pixels, representing the filtered-out background, while the other side comprises non-black pixels denoting the base and honeycomb structure. Leveraging this contrast, we employ convolution with a specially-designed kernel to detect this transition in pixel intensities. This kernel traverses the image as a sliding window, maximizing convolution results precisely at the plastic base boundary. Consequently, this process yields a mask delineating the boundary pixels. As a result, this process generates a mask that marks the boundary pixels. With this mask as a seed, we use morphological operations to generate an enhanced mask that covers the entire plastic base. Once the plastic base is removed, we segment the plastic cell edges. Since there is a distinct difference in intensities between these groups of pixels, we return to histogram filtering, where we identify an optimum threshold value that segments out the peaks corresponding to honeycomb (dark yellow) and plastic (blue), as shown in Fig. S1C. It is worth nothing that the segmentation of plastic cell edges is used solely for visualization purposes (See supplementary movies S1-S7). The final result of our image processing pipeline on the sample with $S = 1$ is depicted in Fig. S1D, showing the segmented 3D volume followed by images of the cross-sections of the data presented in panel A along different orthogonal planes.

#### 3. Analysis of tilt angle

One of the themes observed in our results is the tilting of honeycomb when the initial frames are of size $1 \leq S \leq 2$. To compare and quantify the tilt across various given cell sizes, we measure the angle of tilt of the built cells relative to the plastic base. We achieve this by creating two parallel planes intersecting the honeycomb at the base and the upper layer, shown in Fig. S2A. In each plane we identify the intersecting hexagonal cross-sections (components) corresponding to the hexagonal cavities of individual cells. Fig. S2B shows these components segmented with various colors. Subsequently, we identify the centroids of each of these hexagonal components in both layers, which are highlighted in Fig. S2B, C with red and cyan circles. The components and their centroids are automatically computed by running *Connected Component Analysis* on each plane in the Dragonfly software. Once these contours are established, we calculate the lateral shift in the centroids of hexagons at the upper layer relative to those at the plastic base. In other words, the angle of tilt for each honeycomb cell is computed through simple geometric analysis using the displacement value of the cell centroids from red to cyan, as visualized in Fig. S2C.

### Supplementary Figures

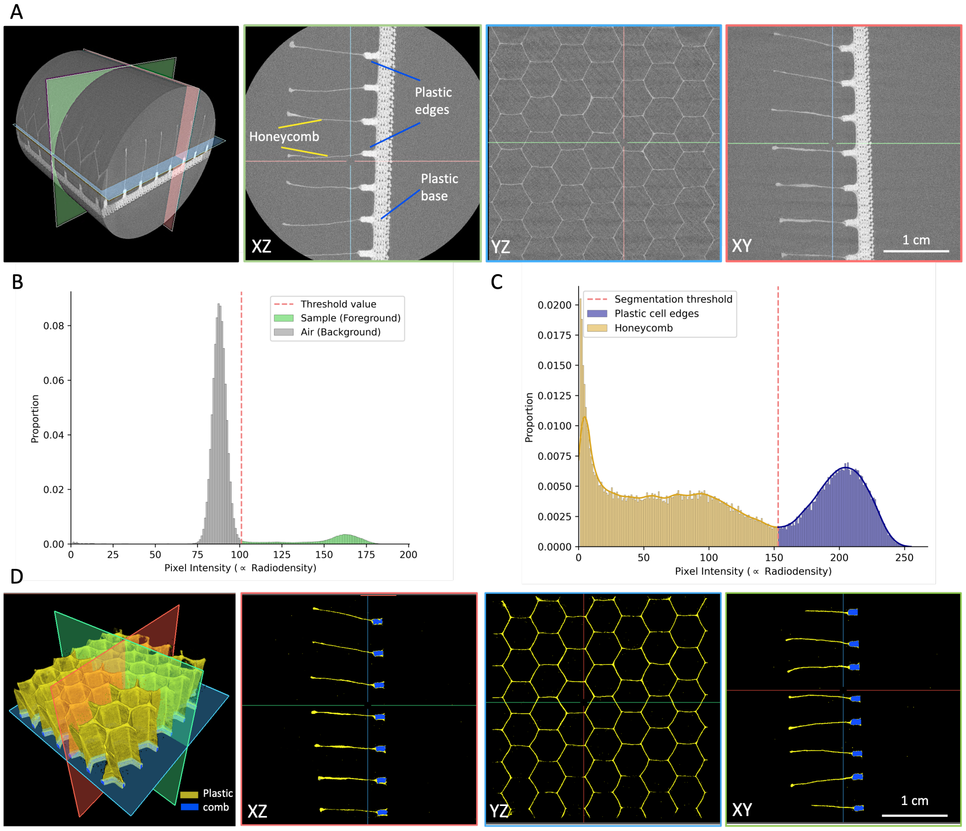


Figure S1: A) X-ray tomography data before processing: the 3D visualization followed by samples of three cross-sections along different orthogonal planes. B) Fraction of pixels with a given intensity in a tomography image showing a sharp peak corresponding to air pixels in the image. The vertical red-dashed line shows the threshold value computed to filter out air (background). C) Histogram thresholding to segment out plastic cell edges. The blue peak corresponds to plastic cell edges and the red-dashed line highlights the optimum threshold value to segment out plastic region. D) Final results after running the complete image processing pipeline, in which honeycomb (dark yellow) and plastic cell edges (blue) are segmented.

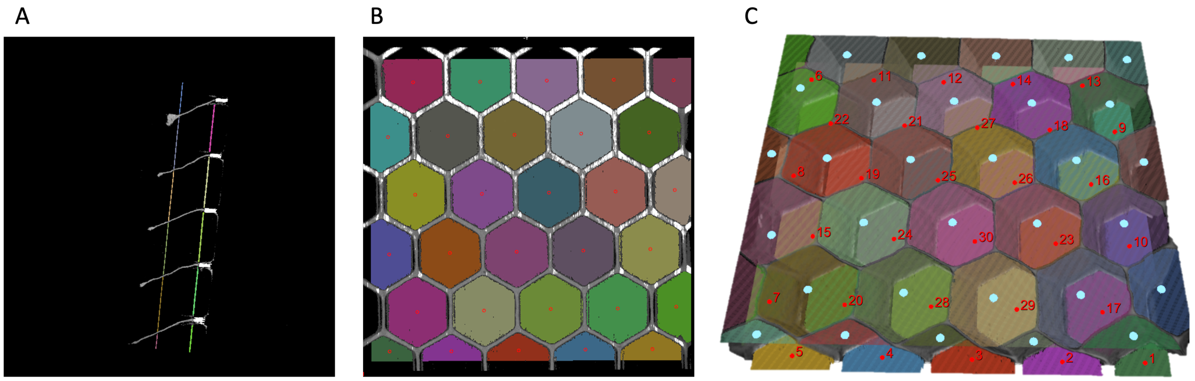


Figure S2: Angle of tilt calculation for individual honeycomb cells demonstrated on a sample with $S = 2$. A) A sample image from the XZ slice, showing the two parallel planes that cross the sample perpendicular to the plastic edges. B) Identifying individual cells using connected component analysis on a cross-section of the sample with the planes shown in A. C) Results of automatic centroid detection on the 3D volume detecting cell centroids in blue (for the top plane) and red (for the lower plane). The angle of tilt for each cell is calculated using the value of the shift in the centroids for each individual cell.

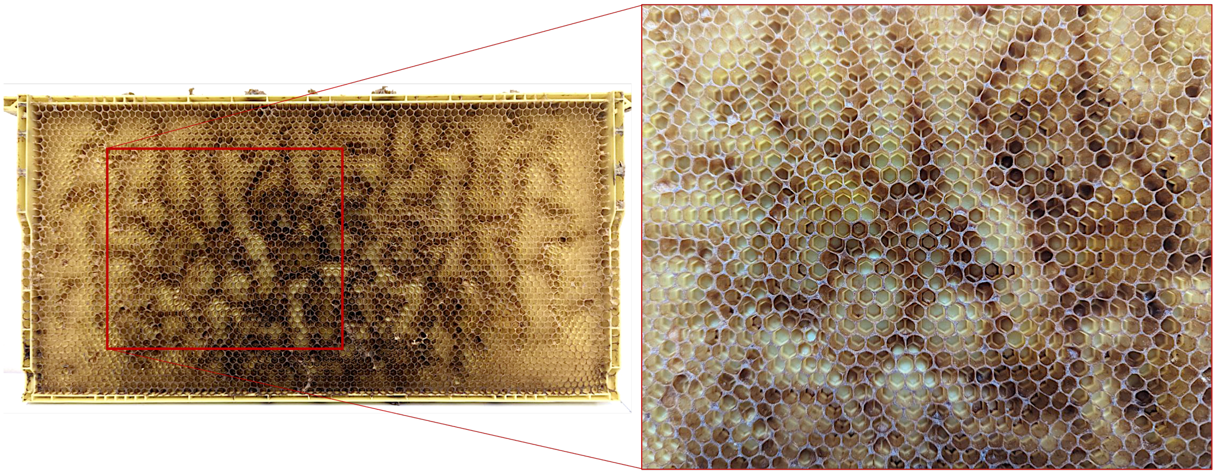

Figure S3: Honeycomb built on one of the commercial frames inside our hives using the merging strategy. A section of comb is magnified to show the occasional combination of the plastic edges on the foundation to build larger cells on top of them.

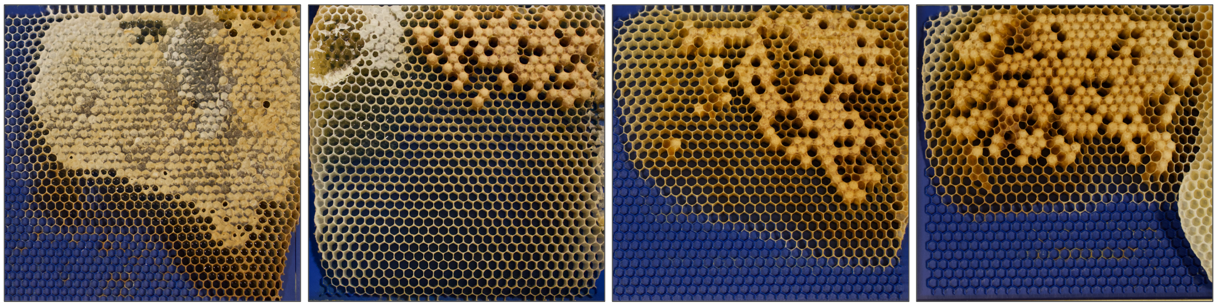

S = 1.25 | S = 1.5 | S = 1.75 | S = 2

Figure S4: Sample frames with larger cell size foundations are either used for raising drone brood or honey storage.

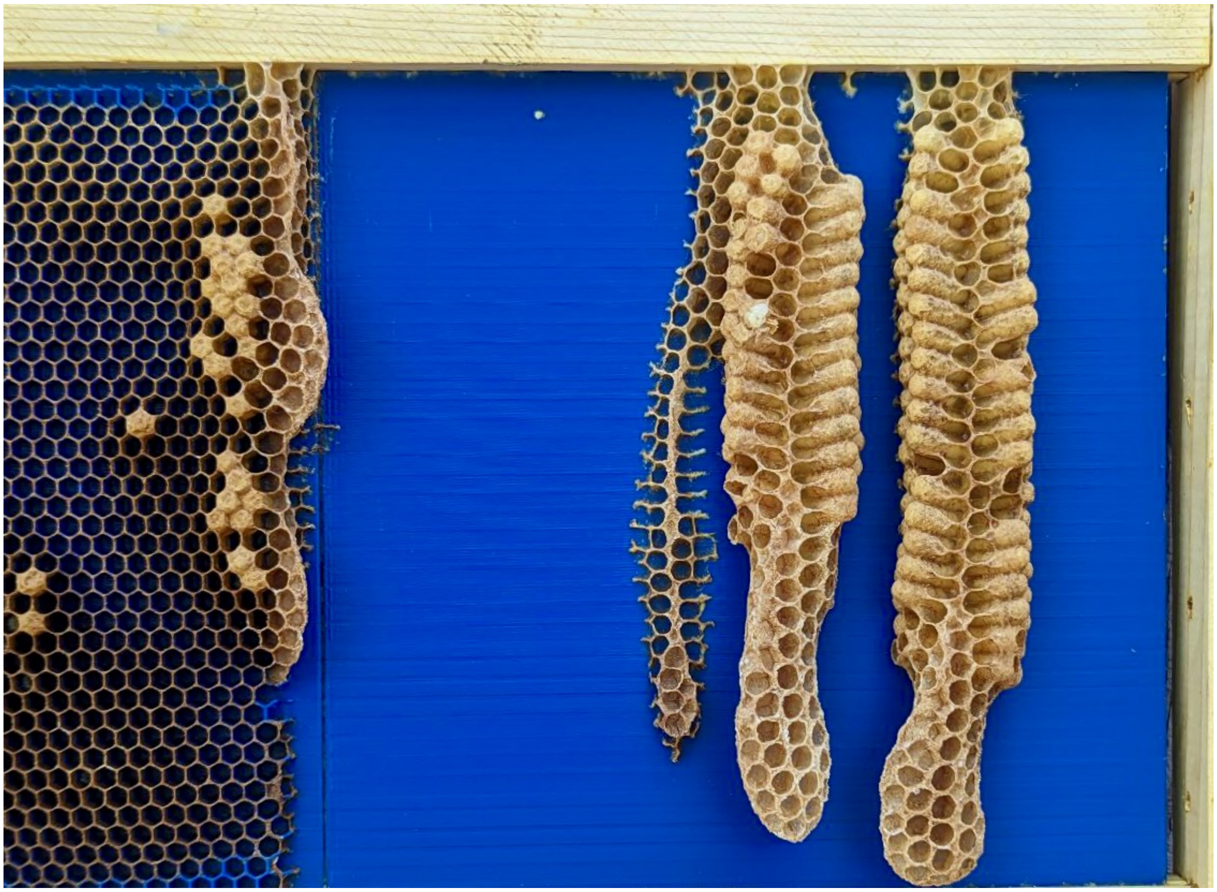

Figure S5: Honeycomb containing drone brood built on the flat side of the experimental frames. This is used to generate the natural tilt angle distribution of the drone comb shown in Fig. 5.

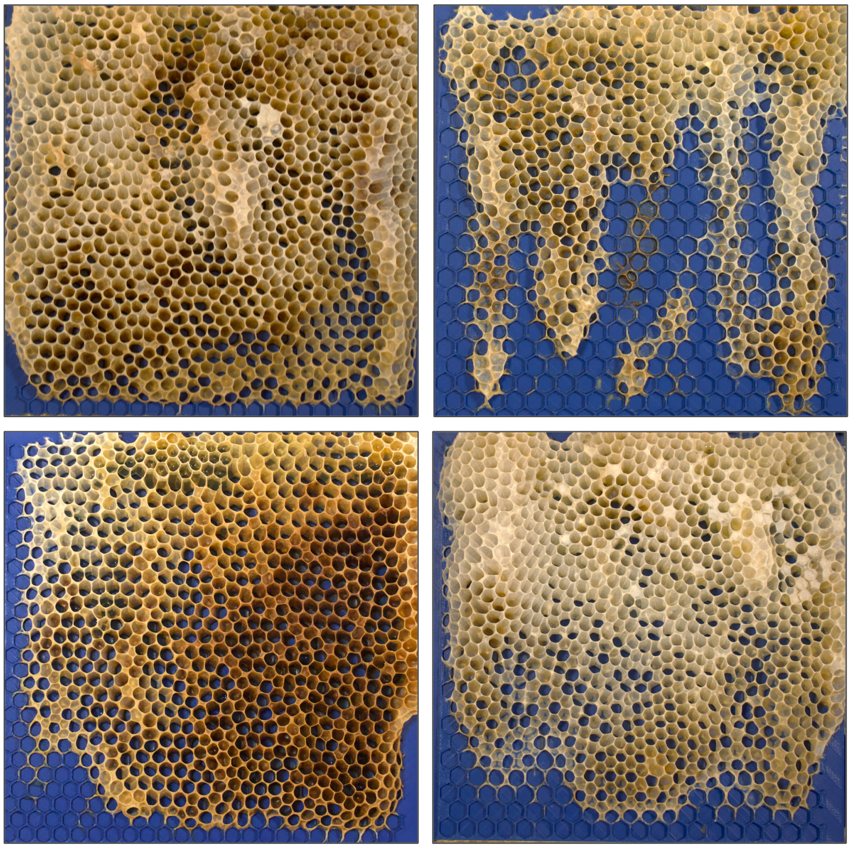

S = 2.5 S = 3.5

Figure S6: Samples of acquired data on frames with $S = 2.5$, and $S = 3.5$ show a combination of tilted or layered modes of building on random sections of the frames.

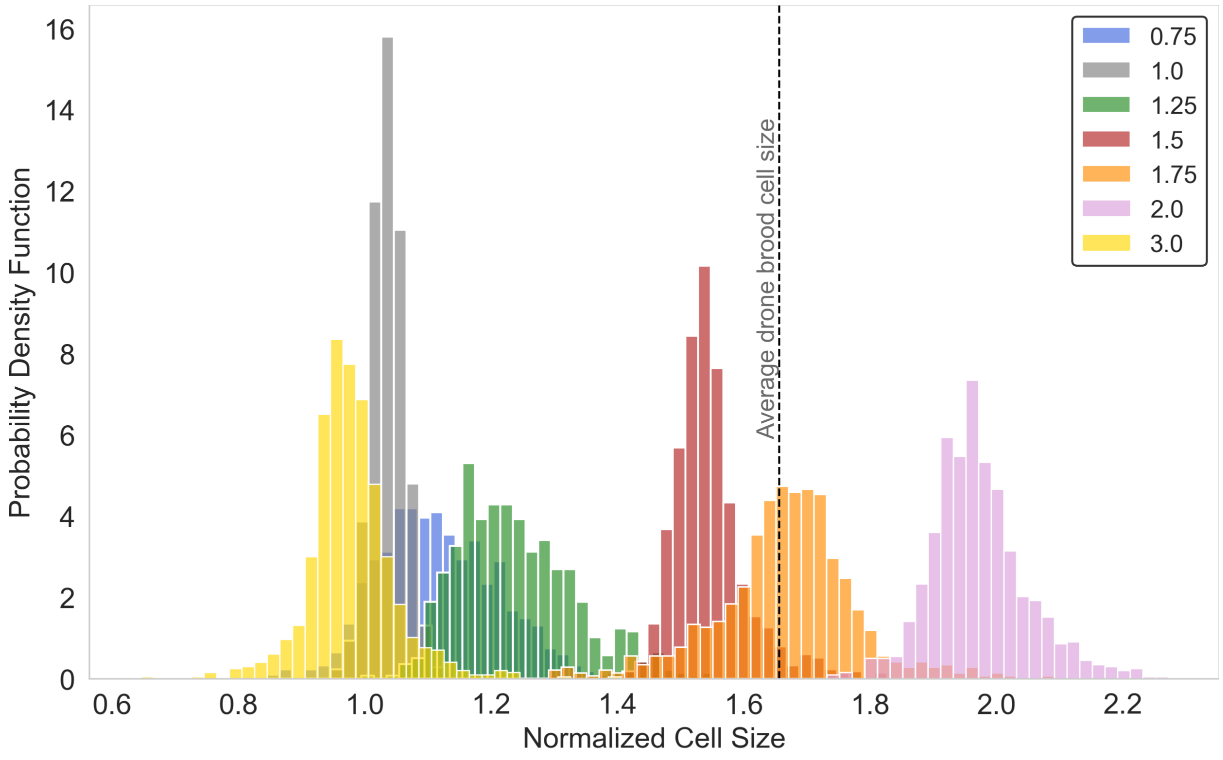


Figure S7: Comparison of cell size distribution of honeycomb built on all of the 3d-printed frames. The given cell sizes ($S$) are shown with different colors in the figure legend.

### Supplementary Movies

Movie S1-S7 show the 3D structure of the honeycomb from various angles, corresponding to the samples with $S = 0.75, 1, 1.25, 1.5, 1.75, 2$, and 3, respectively. These videos can be found in "3D_reconstruction_movies.zip" at `https://datadryad.org/stash/dataset/doi:10.5061/dryad.z8w9ghxmw`. All the 3D volumes are constructed using the X-ray data from the 5 $cm \times 5$ $cm$ sections of the comb constructed on our experimental frames, segmented to show comb in dark yellow and plastic in blue. See the table below for description of the movie files.

| Supplementary Movie | File name | S |
|---|---|---|
| Movie S1 | s_o75_movie.mp4 | 0.75 |
| Movie S2 | s_1_movie.mp4 | 1 |
| Movie S3 | s_1o25_movie.mp4 | 1.25 |
| Movie S4 | s_1o5_movie.mp4 | 1.5 |
| Movie S5 | s_1o75_movie.mp4 | 1.75 |
| Movie S6 | s_2_movie.mp4 | 2 |
| Movie S7 | s_3_movie.mp4 | 3 |

Table S1: Description of the supplementary movie files.

### References


[1] G. Bradski, *Dr. Dobb's Journal of Software Tools* (2000).

[2] Comet Technologies Canada Inc., Montreal, Canada, Dragonfly.

[3] R. C. Gonzalez, R. E. Woods, *Digital image processing* (Prentice Hall, Upper Saddle River, N.J., 2008).
